# Isolation and long-term expansion of a single adult human renal epithelial cell with efficient kidney regeneration capacity

**DOI:** 10.1101/2020.01.21.914762

**Authors:** Yujia Wang, Zixian Zhao, Qiwang Ma, Hao Nie, Yufen Sun, Xiaobei Feng, Ting Zhang, Yu Ma, Jing Nie, Wei Zuo

## Abstract

A wide spectrum of lethal kidney diseases involves the irreversible destruction of the tubular structures which leads to loss of renal function. A reliable technological platform to culture and transplant adult human-derived cells with nephrogenic potential offers great hope to facilitate human kidney regeneration. Here, we show that in an appropriate feeder cell-based culture system, it is feasible to isolate and long-term expand the progenitor-like SOX9+ renal epithelial cells (SOX9+ RECs) from adult mammalians. Single cell-derived SOX9+ REC lines can be established from human needle biopsy or urine samples with molecular homogeneity and genomic stability maintained during culture. Such cells grown in 3D culture system could self-organize into renal organoids composed of proximal tubular, Loop of Henle (LOH) and distal tubular cells as illustrated by single cell transcriptomic analysis. Once being transplanted into the physically injured mouse kidney, the expanded single human SOX9+ RECs incorporated into the damaged area and demonstrated capacity of regenerating functional tubules in vivo. Altogether, the ability to extensively propagate human SOX9+ REC in culture whilst concomitantly maintaining their intrinsic lineage differentiation commitment suggests their future application in regenerative medicine.

---

The isolation and long-term expansion of stem or progenitor cells from adult tissues are fundamental techniques supporting biomedical studies, including disease modeling, drug screening, and cell replacement regenerative medicine. In 1975, Green et al., established the first successful example of adult human epidermal cell culture^1^. Based on a fibroblast feeder system, they were able to culture epidermal stem/progenitor cells as colonies with a remarkable capacity to proliferate. A similar strategy has since been applied to clone epithelial stem/progenitor cells from multiple adult organs and tissues including the cornea, airway, esophagus, and intestine^2–6^. Different from the adult tissue-derived organoid culture system, this feeder cell-based system supports the rapid acquisition of a large number of homologous, primitive cells for the purpose of full characterization and subsequent transplantation, providing a realistic option for cell replacement therapies.

Kidney diseases affect about one in every ten people worldwide and the end-stage renal diseases (ESRD) leads to more than one million deaths globally each year. Most of the lethal chronic kidney diseases (CKD) were characterized by progressive, irreversible destruction of kidney tissue^7^. Besides the mitigating hemodialysis treatment, kidney transplant surgery is so far the only solution for ESRD however its application is quite limited due to lack of donor organ as well as immune rejection^8^. Stem cell-based therapeutic strategies aiming to re-establish the morphology and function of nephron in situ holds great potential to ameliorate or even cure such diseases^9, 10^. Previous studies have shown that pluripotent stem cells (PSC) can be induced into mature renal cells efficiently, offering an attractive cell-source for nephron regeneration^11–13^. For adult kidney, previous studies reported the putative stem/progenitor cell population in adult kidney with nephrogenic potential^14–16^, and other studies demonstrated that the surviving adult epithelial cells after acute kidney injury have the ability to repopulate the damaged nephrons^17, 18^. Altogether the previous investigations have demonstrated that there are distinct populations of nephrogenic cells in kidneys of different developmental stages from embryo to adult, and if such cells can be long-term cultured and properly transplanted into kidney, they should have a broader application in cell therapy and drug screening.

Sex-determining region Y box (SOX) is a family of transcription factors that are widely involved in maintaining cells in a stem/progenitor cell-like state and inhibiting cell differentiation. SOX9 was considered as a marker of stem or progenitor population in some tissues, such as hair bulge, airway epithelium, intestine, pancreas, and neural crest^19–22^. In the normal mouse kidney, rare individual, or small clusters of SOX9+ cells have been reported to be present in the proximal and distal tubules but absent from the LOH or the collecting duct epithelium^23^. In current work, we established an advanced feeder cell-based stem/progenitor cell clone (“F-Clone”) system to isolate and expand nephrogenic SOX9+ REC from adult mammalian kidneys. Utilizing the “F-Clone” system, we achieved long-term, genetically stable expansion of adult human SOX9+ REC, which were isolated from either kidney biopsy or urine. The long-term cultured human SOX9+ REC have the potential to differentiate into distinct types of tubular epithelial cells in 3D culture. After being transplanted into injured immune-deficient mouse kidney, the human SOX9+ REC can incorporate into the kidney and regenerate proximal tubules.

## RESULTS

### Establishing the “F-Clone” system to grow adult SOX9+ RECs from mammalian kidneys

Tissues from the cortex and medulla of normal adult mouse kidneys were dissected respectively and digested to single cell suspensions for cell culture. Firstly we tried conventional culturing method other than “F-Clone” system, which demonstrated very slow tubular cell growth and senescence-like cell morphology (Supplementary Fig. 1a). So we introduced the 3T3 feeder cell based “F-Clone” system plus SCM-6F8 media to selectively isolate and culture clonogenic cells from adult kidney. In F-Clone system, the combination of growth factors and regulators support the maintenance of epithelial tissue stem/progenitor-like cells in a ground-state form as previously reported^3^. Mouse strain constitutively expressing tdTomato protein was used for clonogenic cell isolation. In this culture system, the cells formed compact epithelial colonies and grew up very rapidly (Fig. 1a and Supplementary Movie 1). From 40,000 cells we harvested from kidney cortex, we successfully cultured 18 (±3) cell colonies (n=5 mice) within 4-7 days after seeding, indicating a cell clonogenic ratio of 0.05% approximately. Similar clonogenicity was also noticed in the medulla.

**Figure 1.**
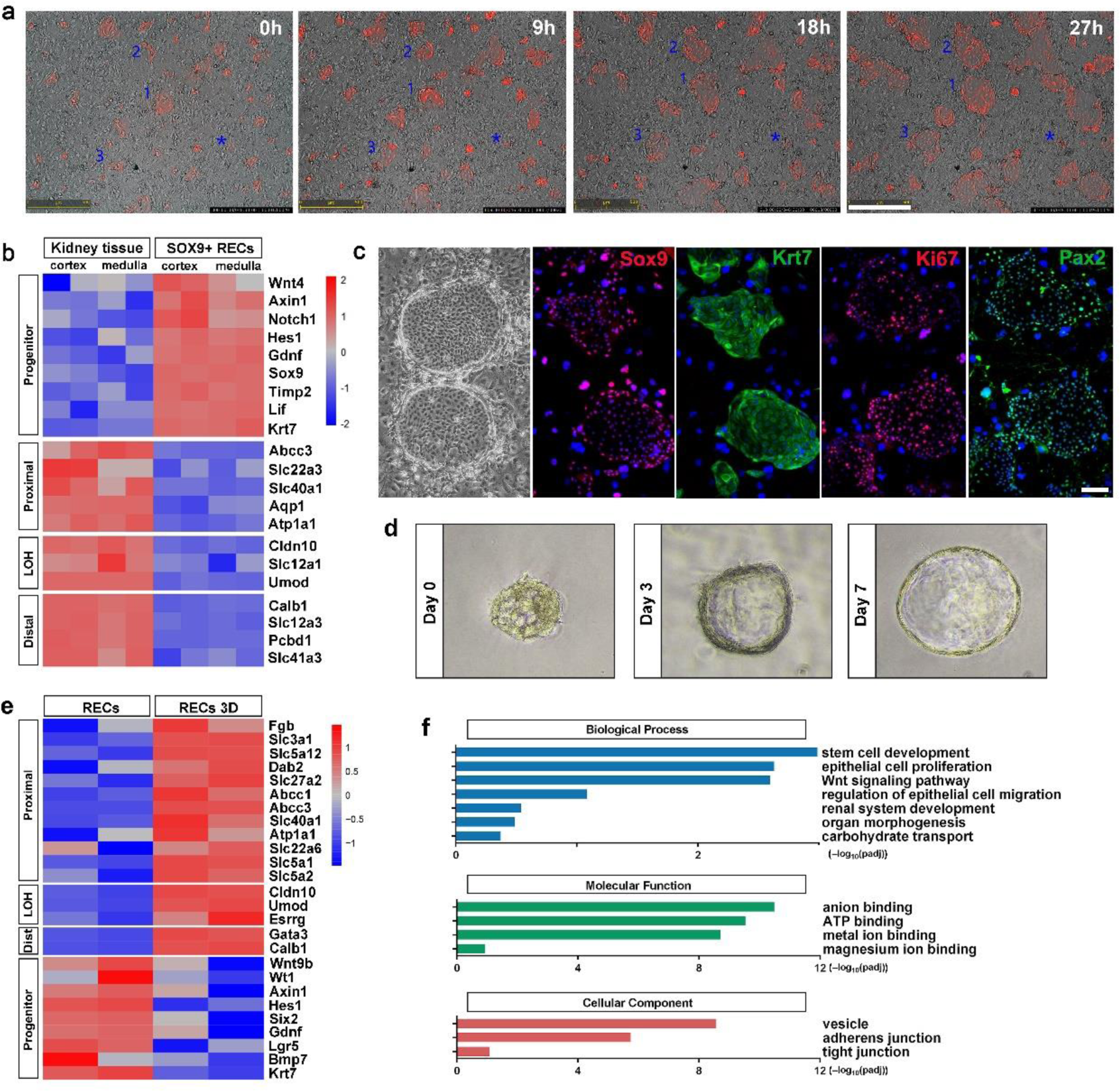
Mouse Sox9+ REC cloning and characterization. (**a**) Representative serial images showing mouse Sox9+ REC single cell-derived clone expansion within 27 hours. Identical visual field were captured by time-lapse microscopic camera. 1-3 indicate the clones with high proliferation capacity. Asterisk indicates the clone with low proliferation capacity. Scale bar, 500 μm. (**b**) Heatmap of differentially expressed gene sets exhibiting a distinct transcriptome between mouse kidney tissue and the corresponding clonogenic Sox9+ REC by microarray analysis. Duplicates were taken from independent biological samples. (**c**) Representative mouse Sox9+ REC colonies immunostained with Sox9, Krt7, Ki67, and Pax2. Scale bar, 100 μm. (**d**) In vitro differentiation of mouse RECs on 3D Matrigel drops showing typical epithelial cell spheres with lumens. (**e**) Heatmap of differentially expressed gene sets of Sox9+ RECs and the cell spheres on 3D Matrigel drops. (**f**) Gene ontology enrichment analysis of genes highly expressed in 3D cell spheres.

In order to characterize the clonogenic cells derived from the cortex and medulla, transcriptome microarray analysis was performed with the clonogenic cells and their corresponding kidney tissues. Compared to normal kidney tissue, the cells derived from both locations displayed high expression of Sox9 and also a number of genes previously known as mediators of prenatal renal development such as Gdnf, Lif, and Krt7. In contrast, the clonogenic cells did not express the marker genes of the mature renal cells (Fig. 1b and Supplementary Fig. 1b). Gene ontology analysis also presented an extensive enrichment of genes linked to kidney development and cell proliferation in progenitors (Supplementary Fig. 1c). Immunostaining confirmed that all the single cell-derived clones are positive for Sox9 and Krt7, and also stained positive for proliferative marker Ki67 and embryonic renal marker Pax2 (Fig. 1c), while negative for those mature renal epithelium markers (Supplementary Fig. 2).

Next we evaluated whether the mouse clonogenic Sox9+ renal epithelial cells (RECs) have lineage differentiation potential in vitro. Sox9+ RECs were grown in the solidified reconstituted basement membrane (Matrigel matrix) -based 3D culture system for differentiation. Cells gradually self-organized into sphere structures within 7∼12 days (Fig. 1d). Transcriptomic analysis in Fig. 1e showed that comparing to the undifferentiated SOX9+ RECs, the 3D spheres were characterized by under-expression of multiple progenitor-related genes such as Six2, Krt7 and Lgr5 while over-expression of multiple tubular markers including those of the proximal tubule (Atp1a1/Slc5a1/Slc5a2), LOH (Umod), and distal tubule (Calbindin). Gene ontology analysis showed enrichment of related biological processes including “renal system development”, “regulation of epithelial cell migration” and “organ morphogenesis” in 3D spheres. Enriched molecular functions included anion binding and metal ion binding, which are required for the ion transport function of renal tubules (Fig. 1f).

Previous study showed that Sox9+ RECs are very rare in healthy mouse kidney but can be activated in various acute kidney injuries (AKI) mouse models^23, 24^. Here we introduced a new AKI model which involves unilateral partial nephrectomy (UPN) to characterize and isolate Sox9+ REC. In this UPN model, about one third of the renal tissue were removed by surgery, which involves both cortex and medulla. One day post-UPN surgery (1 dps), we observed strong expression of kidney injury marker Kim-1^25^ along the cut edge of the cortex, but not medulla. Simultaneous emergence of a large number of Sox9+ RECs was observed surrounding Kim-1 positive tubular area (Fig. 2a-c). The newly appeared Sox9+ RECs had not yet entered cell cycle on 1 dps as only 1.6(±0.5)% of them stained positive for Ki67 antigen. However, the cells became further activated afterward as the Ki67+ cell ratio increased to 18.3 (±9.9)% on 3 dps and 47.6 (±9.2)% on 7 dps. In contrast, most of the endogenous Sox9+ cells in uninjured healthy kidney were in quiescent status as only 3.7(±2.6)% of them stained positive for proliferative marker Ki67. Our observation was consistent with previous studies which have demonstrated that the endogenous Sox9+ cell activation and expansion is an early event after acute renal injury^23, 24^.

**Figure 2.**
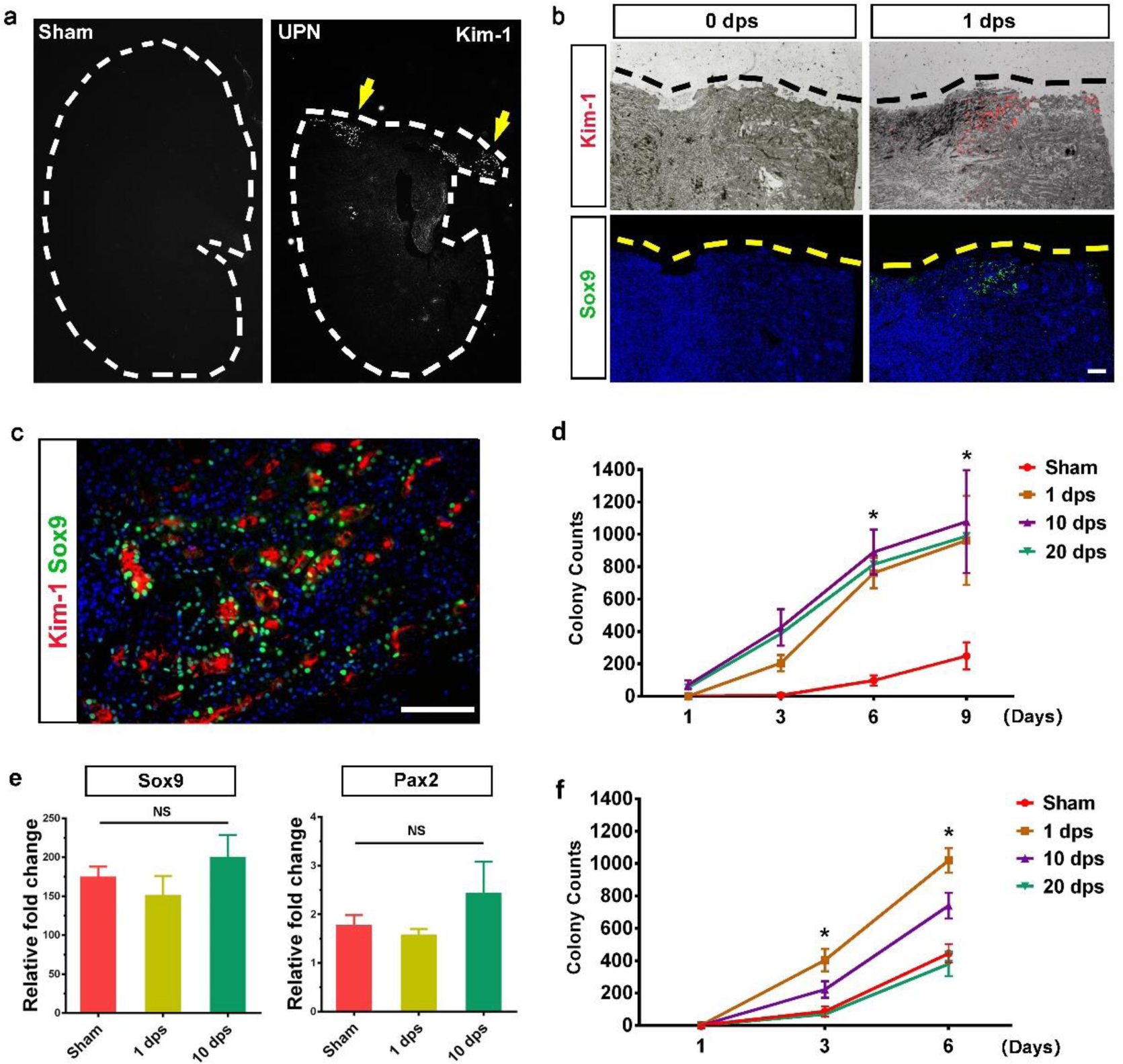
Sox9+ REC activation in acutely injured kidneys. (**a**) Immunostaining of kidney injury marker Kim-1 (grey) in the UPN injured kidney. The mouse kidney was analyzed 24 hours after partial nephrectomy on the upper 1/3 part of left kidney. Yellow arrows indicate Kim-1+ cells by immunostaining. (**b** and **c**) Co-staining of Kim-1 and Sox9 along the cutting edge of the injured kidney. Black and yellow dashed lines indicate the blade cut edge. Scale bar, 500 μm. (**d**) Growth curve of the colonies derived from sham and UPN injured kidneys (n=6 independent mice in each group). The whole left kidney was analyzed for clonogenic assay at indicated days after surgery. Statistics are inclusive of all biological replicates. *p < 0.05. (**e**) qRT-PCR analysis of transcripts of Sox9 and Pax2 in RECs derived from sham and partial nephrectomy injured kidneys (n = 6 independent qRT-PCR experiments). Statistics are inclusive of all biological replicates. NS, no significant difference. (**f**) Growth curve of the colonies derived from sham and UUO injured kidneys (n=6 independent mice in each group). The whole left kidney was analyzed for clonogenic assay at indicated days after obstruction surgery. Statistics are inclusive of all biological replicates. *p < 0.05.

Then we collected the whole UPN kidney and grow the cells using the F-Clone protocol. As expected, we isolated larger number of Sox9+ RECs from UPN kidneys than from the healthy kidneys, and the 1 dps group had the highest clonogenicity (Fig. 2d). The Sox9 and Pax2 expression levels in Sox9+ REC from UPN kidneys were almost identical to those from healthy kidneys, suggesting that they share identical molecular features (Fig. 2e and Supplementary Fig. 3a). Similar clonogenic results were obtained from another unilateral ureteral obstruction (UUO) kidney injury model^26^ (Fig. 2f and Supplementary Fig. 3b). Therefore, our “F-Clone” system can be used to grow progenitor-like Sox9+ RECs from both healthy and acutely injured kidneys.

To confirm the applicability of the “F-Clone” system in other higher animals, we also successfully cloned Sox9+ REC from cortex, medulla, and papilla of adult rhesus monkeys using the similar protocol (Supplementary Fig. 4).

### Characterization of long-term cultured SOX9+ REC derived from adult human kidney and urine

Previous studies suggested that the progenitor-like cells identified in rodent systems may not formally extrapolate the situation in the human kidney^27–29^. Therefore our successful isolation of REC from the injured mouse kidney encouraged us to further test the “F-Clone” strategy to grow SOX9+ REC in human patients. Single-cell suspensions of needle biopsy tissue derived from three patients with CKD were seeded for cell culture and all of the samples successfully yielded cell colonies within 3∼7 days. Such human cell colonies grown on mouse feeder cells express human-specific nuclei antigen and a great part of them express proliferative marker Ki67 (Fig. 3a). And as renal biopsy is an invasive procedure with high risk for patients, we also seek other solution to isolate the clonogenic cells. For CKD patients who were not eligible for biopsy, or for healthy adults, we cloned cells from their urine. We tested three healthy adult urine samples and successfully obtained clonogenic cells from two (66.6%) of them. We also tested seven CKD patient urine samples and successfully obtained clonogenic cells from five (71.4%) of them. Averagely every 10,000 adult human kidney or urine-derived cells seeded on feeder cells will yield about one cell colony. All adult human sample information is listed in Supplementary Table 1.

**Figure 3.**
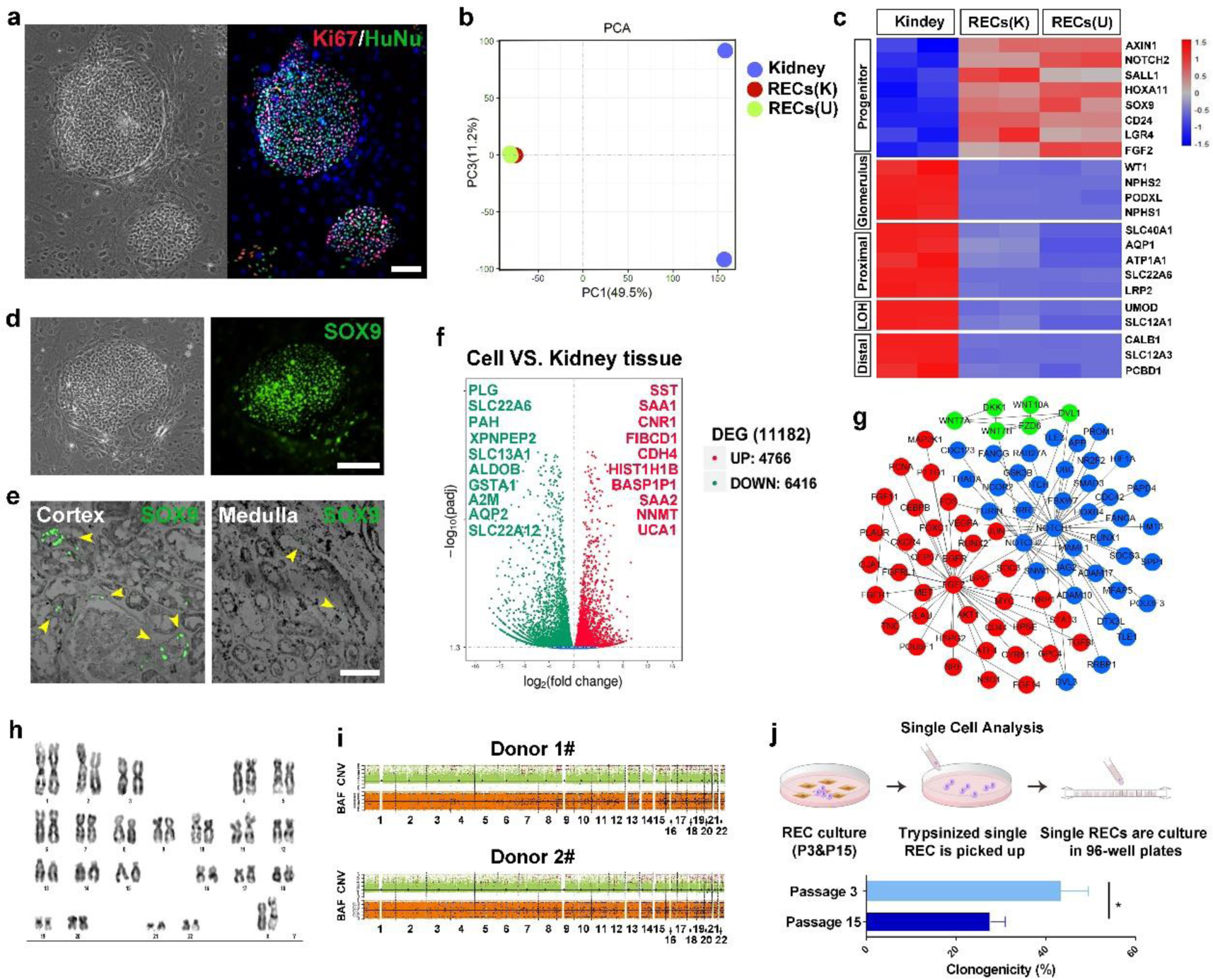
Cloning of adult human SOX9+ REC from kidney biopsy and urine. (**a**) Human renal epithelial cell colonies from kidneys of patients with CKD stained with proliferative marker Ki67 and human-specific marker human nuclear antigen HuNu (representative image of n=3 independent human cell pools). Scale bar, 100 μm. (**b**) PCA map showing whole transcriptomic profiles of kidney tissue, kidney-derived SOX9+ RECs (K) and urine-derived SOX9+ RECs (U). Duplicates were taken from independent biological samples. (**c**) Heatmap of differentially expressed gene sets of human kidney tissue and the corresponding clonogenic SOX9+ REC by RNA-seq analysis. Duplicates were taken from independent biological samples. (**d**) Human SOX9+ REC colonies immunostained with SOX9 antibody. Scale bar, 100 μm. (**e**) Endogenous SOX9 expressing cells in human cortex and medulla tissue samples. Yellow arrowheads indicate SOX9 expressing cells. Scale bar, 50 μm. (**f**) Volcano plot of up-regulated (red) and down-regulated (green) gene expression showing significant differences in human kidney-derived SOX9+ REC versus human kidney tissue with top-ranked genes listed. (**g**) Protein-protein interaction network of selected genes with high expression levels in SOX9+ REC clones versus human kidney tissue. (**h**) Representative karyotype image of human SOX9+ RECs after 25 passages. n=3 independent experiments. (**i**) CNV and BAF profiles of 2 human SOX9+ REC lines at passage 14. Regions of copy number gain/loss and loss of heterozygosity regions were shown. Control-FREEC methodology was used to discerns somatic variants from germline ones. Predicted BAF and copy number profiles are shown in black. Gains, losses and loss of heterozygosity are shown in red, blue and light blue dots, respectively. (**j**) Scheme demonstrating the single cell picking by 96-well plates followed by quantification of single cell clonogenicity at early (passage 3) and late (passage 15) passage of human SOX9+ REC. n = 3 individual clonogenic assays using independent biological samples.

The clonogenic cells isolated from renal biopsy and urine are indistinguishable in either morphology or growth kinetics. RNA-Seq analysis showed that they shared great similarity at whole transcriptome level with high expression progenitor markers including SOX9, SALL1, HOXA11 and LGR4 (Fig. 3b and 3c). Immunostaining confirmed the SOX9 expression in the cell colonies of healthy adults and patients (Fig. 3d and Supplementary Fig. 5a), suggesting the clonogenic human cells were the counterpart of SOX9+ RECs in the mouse kidney. Indeed, we observed endogenous SOX9+ REC in human kidney tissue samples and there were nearly 5-fold more endogenous SOX9+ REC in cortex than in the medulla (Fig. 3e).

The cultured human SOX9+ REC did not express the mature tubular or glomerular markers by RNA-Seq or immunostaining (Fig. 3c and Supplementary Fig. 5c). By RNA-Seq analysis, we identified a number of novel progenitor cell markers as listed in Fig. 3f. Among them, two serum amyloid A genes (SAA1 and SAA2) were listed in the highly expressed gene list, and indeed the SAA gene was previously reported to be able to associate with regenerative capacity of kidney^30^. Novel cell surface markers such as SST, CNR1, and CDH4 were also identified which could be used to flow sorting of SOX9+ REC. A few critical signaling pathway core members were also highly expressed in cultured SOX9+ REC including NOTCH1/NOTCH2/JAG2, Wnt7A/Wnt7B/Wnt10/FZD6/DVL1, and FGF2/FGF11/FGF14/FGFR1. NOTCH, Wnt, and FGF pathways were all known to play a pivotal role for kidney development and pathology^31–36^ and we found that they worked in a protein-protein interaction network to maintain the growth of SOX9+ REC in culture (Fig. 3g).

The human SOX9+ REC cultures can be stably maintained for long-term expansion. One line in our lab has been passaged for six months and 25 generations, which could yield approximately 1x10^20^ cells with normal karyotype maintained (Fig. 3h). Also, whole-genome copy number variation (CNV) profiling^37, 38^ revealed a very limited spectrum of somatic chromosomal copy number changes existed in the long-term cultured RECs (Fig. 3i).

In order to characterize the long-term propagation ability at single cell resolution, we manually picked 100 single cells from early (Passage 3) and late (Passage 15) passages to compare their clonogenic capacity. Single cells cloned from early passage are morphologically identical to the late passage ones. However, during subsequent clonal plating, the average clonogenicity of cells decreased from 42% (early passages) to 23% (late passages) (Fig. 3j). No elevation of mature tubular or glomerular markers was observed during passage (Supplementary Fig. 5b). This result confirmed the transit-amplifying progenitor-like properties of the cells that could replicate for long-term without spontaneous differentiation yet with gradual loss of clonogenicity^39^.

Next we evaluated whether the progenitor-like human SOX9+ RECs have lineage differentiation potential in vitro. Previous studies on human iPS cells demonstrated efficient kidney eorganoid formation in 3D Matrigel culture system^40, 41^. In our experiments, SOX9+ RECs in 3D Matrigel culture system formed spheres with lumen structures in 7∼12 days (Fig. 4a). RNA-Seq transcriptomic analysis showed that the Matrigel spheres were characterized by under-expression of multiple progenitor genes such as LGR4 and SALL1 and over-expression of multiple tubular markers including those of the proximal tubule (ATP1A1/SLC22A6/AQP1), LOH (SLC12A1), and distal tubule (SLC12A3) (Fig. 4b). No glomerular lineage marker expression was detected in such Matrigel spheres.

**Figure 4.**
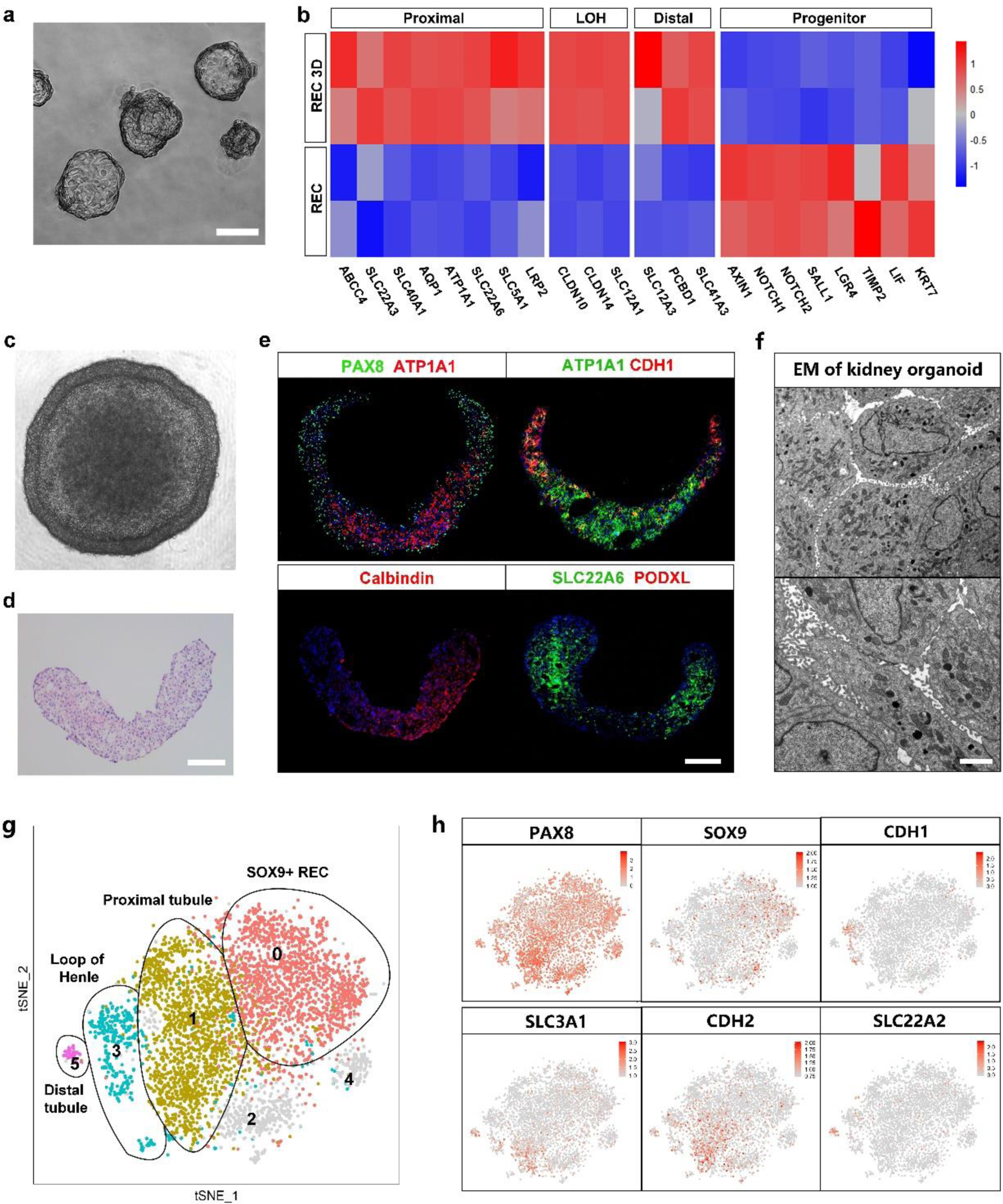
In vitro 3D differentiation of human SOX9+ REC. (**a**) Representative image of cell spheres of adult human single cell-derived SOX9+ REC lines in 3D Matrigel culture system. Scale bar, 50 μm. (**b**) Heatmap of the gene sets differentially expressed between single cell-derived SOX9+ REC and the corresponding SOX9+ REC 3D cell spheres. Duplicates were taken from independent biological samples. (**c**) Suspension 3D culture of adult human single cell-derived SOX9+ REC. Scale bar, 100 μm. (**d**) Histology of single cell-derived SOX9+ REC organoids in suspension culture after longitudinal sectioning. Scale bar, 100 μm. (**e**) Immunofluorescence on longitudinal sections of 3D organoids in suspension culture with indicated tubule and podocyte markers. Scale bar, 100 μm. (**f**) Representative TEM of human single cell-derived SOX9+ REC organoids in suspension culture showing characteristic tubule morphology. Scale bar, 2 μm. (**g**) Unsupervised graph-based clustering identifying 6 transcriptionally distinct cell clusters in single cell-derived SOX9+ REC organoids. (**h**) Gene expression levels of PAX8 (pan-tubule), SOX9 (REC), CDH1 (LOH), SLC3A1 (proximal tubule), CDH2 (proximal tubule), and SLC22A2 (distal tubule) demonstrated by a tSNE featureplot.

### Human SOX9+ RECs in 3D suspension culture self-organized into kidney organoids

Previous studies on human pluripotent stem cells also demonstrated efficient kidney organoid formation in specialized 3D suspension culture system. In our experiments, 96-well ultra-low attachment plates with the U-shaped bottom were used to support the 3D suspension culture of SOX9+ REC and facilitate the formation of tubular aggregates. After 12∼14 days culture, SOX9+ RECs self-organized into organoids which recapitulated the characteristic structure of kidney with one convex border and another concave border as shown by longitudinal sectioning & staining (Fig. 4c and 4d). Immunostaining in Fig. 4e revealed expression of tubular epithelium markers in the organoids including the PAX8 (pan-tubule), ATP1A1 (proximal tubule), SLC22A6 (proximal tubule), CDH1 (LOH) and Calbindin (distal tubule). Transmission electron microscopy (TEM) also confirmed the characteristic proximal tubular cell features such as brush border microvilli and enriched mitochondria in cytoplasm (Fig. 4f).

To illustrate the cell composition of the kidney organoids, 10X genomics single-cell RNA sequencing was performed. We isolated and sequenced 5,848 cells, and eventually analyzed 5,098 cells after stringent quality control. We performed unsupervised graph-based clustering (Seurat method) of the dataset and identified 6 transcriptionally distinct cell clusters (Fig. 4g), which were visualized by t-distributed stochastic neighbor embedding (tSNE). Most of the organoid cells were PAX8+ with very few cells expression glomerular cell markers, confirming their tubular epithelium cell identity. Based on known cell type-specific markers, four major clusters (Cluster 0, 1, 3, 5) were annotated as undifferentiated RECs (SOX9+)，proximal tubular cells (SLC3A1+/CDH2+), LOH cells (CDH1+) and distal tubular cells (SLC22A2+), respectively (Fig 4h). The identity of other 2 clusters remained unclear. Cell number counting indicated that 37.2% cells were proximal tubular cells, 9.3% cells were LOH cells, while merely 1.0% were distal tubular cells. It seems that the differentiation fate commitment of human SOX9+ RECs is biased to the more proximal cells in the tubular system. Altogether, the data above demonstrated that the long-term cultured human SOX9+ RECs had differentiation potential to give rise to mature tubular epithelial cells in vitro.

### Precondition mouse kidney for intra-renal transplantation by wedge resection surgery

Next in order to test the in vivo regenerative capacity of long-term cultured human SOX9+ REC, we labeled the single cell-derived SOX9+ REC with GFP-expressing retroviral vector and tried to transplant the cells to injured kidney immune-deficient NOD-SCID mice. First, we tried delivering the GFP-labeled SOX9+ RECs via tail vein injection into the injured mouse kidney. However, the results showed no engraftment of human cells in the mouse kidneys regardless of the injury model used (IRI, UUO or UPN). In fact, most the injected human cells were found trapped in blood vessels of mouse lungs. To bypass the capillary barrier, we also tried injecting human cells into renal artery but still observed no engraftment (data not shown).

Then we tested whether the SOX9+ RECs can be directly injected into the mouse kidneys to achieve engraftment. When we directly injected the GFP+ cells into the parenchyma of healthy kidney, it led to minimal GFP+ cell distribution beneath the surface renal capsule and no engraft was observed inside the kidney. Then we realized that preconditioning the kidney with some kind of physical injury (eg.incision) would be necessary for successful intra-renal transplantation. Such a physical injury would not only make an intra-renal space but also create an inflammatory environment for cells to attach and grow. To this end, a wedge resection surgery were performed on one kidney of mice. Most mice survived well after the manipulation but lost significant body weight because of the stress. H&E staining showed massive tissue injury, large-scale immune cell infiltration and size shrink of kidney (Fig. 5a). Their serum creatinine level had a sharp rise one day after surgery and gradually recovered after 1∼3 weeks, but was still approximately 2-fold higher than the baseline (Fig. 5b).

**Figure 5.**
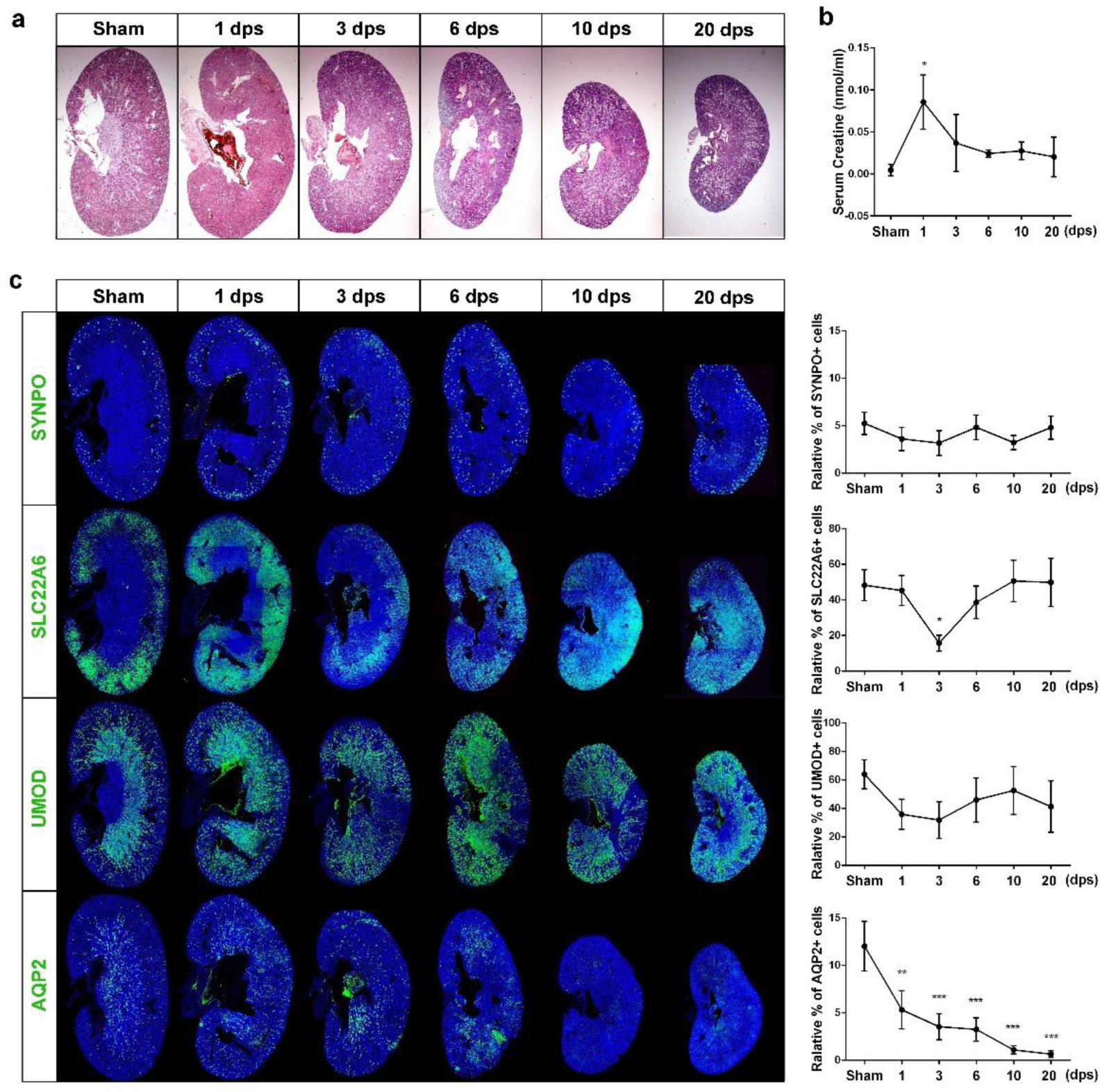
Wedge resection model that induces acute kidney injury. (**a**) Representative images of H&E stained whole kidney sections at indicated time points. dps, days post surgery. n=5 individual injury experiments using independent biological samples. (**b**) Serum creatinine level measurement in sham and injured mice. dps, days post surgery. n=5 individual injury experiments using independent biological samples. * P<0.05 compared to Sham control. (**c**) Representative images and quantification of the immunostained whole kidneys harvested at indicated time points. The expression pattern and quantification of tubular and glomerulus markers indicated partially restoration of structures. dps, days post surgery. n=5 individual injury experiments using independent biological samples. * P<0.05, ** P<0.01, *** P<0.001 compared to Sham control.

We further evaluated the tubular damage level after wedge resection. The AQP2+ collecting ducts were permanently damaged by incision. The SLC22A6+ proximal tubule and UMOD+ LOH/distal tubules were largely damaged within first 3 days, which were then partially repaired afterwards. The SYNPO+ glomerulus were free of damage (Fig. 5c). Then we will use this novel preconditioning model to facilitate intra-renal cell transplantation.

### Single cell-derived SOX9+ REC regenerated human kidney tubules in vivo

Next we sorted a single cell-derived SOX9+ REC line labeled with GFP, expanded it to 3x10^6^, and then transplanted the cells into the incision of injured kidney. Transplanted kidneys were analyzed 1∼2 weeks after transplantation. Large-scale engraftment of fluorescent cells into the mouse kidney was observed (Fig. 6a). Frozen sectioning and immunostaining demonstrated incorporation of GFP+ cuboidal epithelial cells into the incised part which lines structures with lumens in the mouse kidney). All GFP+ cells express human-specific nuclei antigen, which proved that the fluorescent tissues were originated from human tissue but not any technical artifact such as autofluorescence (Fig. 6b). Approximately 3.7% (±1.1%) of the engrafted human cells expressed proliferative marker Ki67 (Supplementary Fig. 6a).

**Figure 6.**
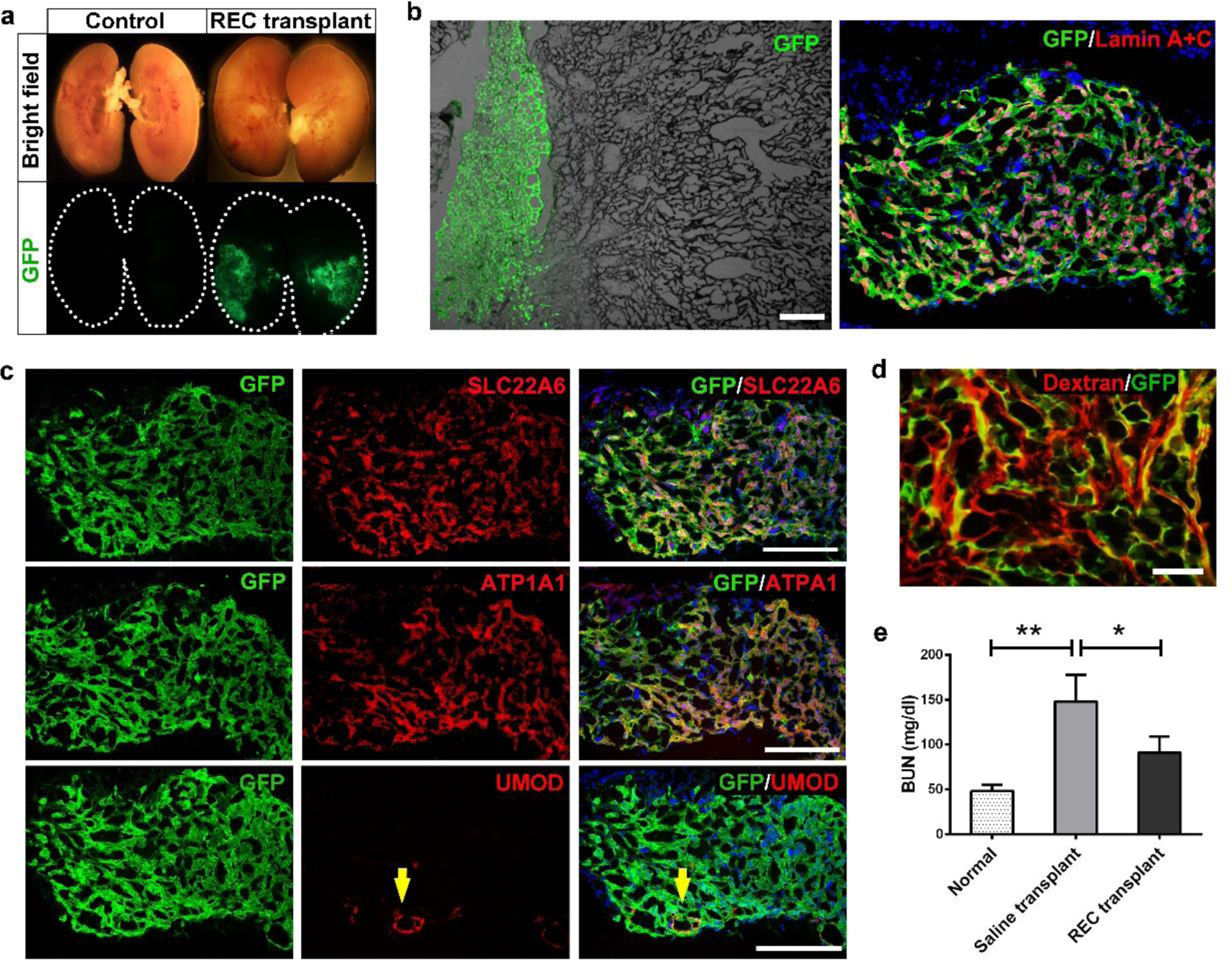
Intrarenal transplantation of human single cell-derived SOX9+ REC. (**a**) Bright field and direct fluorescence of injured NOD-SCID mouse kidney 14 days after saline or single cell-derived GFP+ SOX9+ REC intrarenal transplantation. Kidneys were longitudinally cut open in the midline for direct observation of GFP signal. n=10 independent transplantation experiments. (**b**) Representative bright field and immunostaining images on frozen sections of the transplanted kidney 14 days after transplantation. The descendants of GFP+ single cell-derived SOX9+ REC demonstrated lumen structures. Scale bar, 50 μm. Lamin A+C, a human specific nuclei antigen. (**c**) Frozen sections of transplanted kidney 14 days after single cell-derived SOX9+ REC transplantation followed by immunostaining with GFP and indicated tubule markers. Yellow arrows indicate UMOD+ GFP+ cells by immunostaining. Scale bars, 100 μm. (**d**) Rhodamine-Dextran uptake assay on the regenerated GFP+ human tubules. Scale bar, 50 μm. (**e**) BUN level measurement in normal and injured kidneys with saline or human REC transplantation. n = 6 independent injury and transplantation experiments. * P<0.05, ** P<0.01. All data are presented as mean ± SD.

All of the GFP+ cells express pan-tubular epithelial cell marker PAX8 (Supplementary Fig. 6a), and most of them expressed proximal tubular cell marker genes SLC22A6 and ATP1A1 (Fig. 6c). Meanwhile, a small number of GFP+ cells (<1%) form lumen structures and express UMOD, which is a marker gene mainly expressed in ascending limb of LOH and occasionally the early distal convoluted tubules (Fig. 6c).This biased differentiation fate proportion in vivo is consistent with the 3D organoid differentiation data above. Of note, the morphology of the lumen structure formed by transplanted human cells were still different from native mouse tubules, suggesting that the mouse kidney microenvironment cannot support complete maturation of the human SOX9+ RECs, probably due to insufficient cross-reactivity of multiple growth factors or cytokines between rodent and human.

Dextran uptake assay revealed the accumulation of fluorescent low-molecule-weight dextran in GFP+ human tubules, which suggested that at least some of the regenerated proximal tubular cells could be connected to glomerular filtrate (Fig. 6d). Further blood analysis demonstrated that the regenerated tubules contributed to the recovery of mouse renal function after the injury as shown by the decrease of blood urine nitrogen (BUN) level after REC transplantation (Fig.6e).

The proximal tubules represent a particular target for nephrotoxicity and cisplatin is one such nephrotoxicant which could induce apoptosis of proximal tubular cells. When cisplatin were intraperitoneally given to the transplanted mice (Fig. 7a), we observed significant apoptosis of the GFP+ human tubular cells. Meanwhile, the GFP+ human lung-derived SOX9+ basal cells and their progeny engrafted in lung were unaffected by the cisplatin, suggesting the nephrotoxicity specificity (Fig. 7b and 7c). Altogether the data above demonstrated that a single human SOX9+ REC can regenerate functional tubules in vivo.

**Figure 7.**
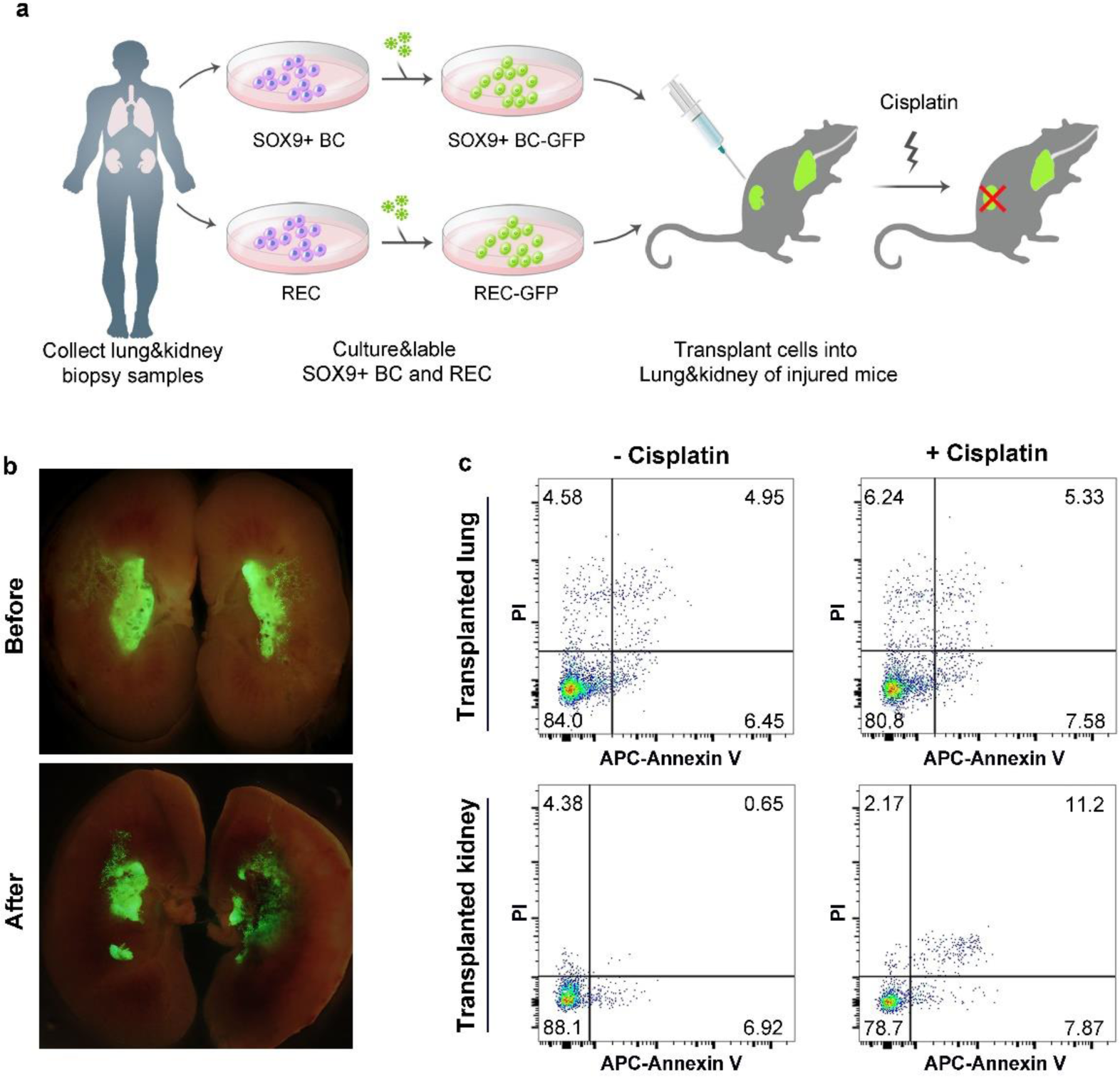
Regenerated human proximal tubular cells were sensitive to cisplatin toxicity. (**a**) Schematic illustrating the generation of dual transplanted mice and the following cisplatin treatment. Human lung-derived SOX9+ basal cells were transplanted into NOD-SCID mouse lung; human kidney-derived SOX9+ cells were transplanted into NOD-SCID mouse kidney. 11 days after the double orthotopic transplantation, 10 mg/kg cisplatin was injected to dual transplanted mice subcutaneously. 3 days after cisplatin treatment, mouse lungs and kidneys were harvested for analysis. BC, basal cell. (**b**) Direct fluorescence on transplanted kidneys showing the transplanted human cells with or without cisplatin treatment. (**c**) Flow cytometry apoptosis analysis on the GFP+ cells of cisplatin treated dual transplanted mice as assessed by APC-Annexin V and propidium iodide (PI) staining. n=3 independent experiments in each group.

In order to investigate the organ-specificity of SOX9+ REC mediated regeneration, we compared the kidney-derived SOX9+ REC with human lung-derived SOX9+ basal cells^42^. We transplanted equivalent number of lung-derived SOX9+ REC into the mouse kidney and found only minimal incorporation. Therefore, the successful engraft of SOX9+ REC into the kidney seems to be an organ-specific process (Supplementary Fig. 6b and 6c).

## Discussion

In this study, we report the robust “F-clone” system that allows selective cloning, long-term proliferative expansion of SOX9+ RECs derived from healthy and diseased human. This work provides the first glimpse into the characteristics of the human SOX9+ REC and reveal their tubular epithelium differentiation potential. Our observation is in line with previous studies in mouse AKI model which revealed that activated Sox9+ progenitor cells were capable to repair the renal tissue, suggesting an evolutionary conserved molecular feature of SOX9+ cells in mammalian^23, 24^. In mouse, most of the AKI-activated Sox9+ REC are derived from expansion/dedifferentiation of survived tubular cells^17, 43–46^, and now in our clonogenic assay, we showed that such activated Sox9+ RECs were similar to the rare Sox9+ cells in healthy kidney.

Previous studies suggested the existence for adult human kidney stem/progenitor cells^47–54^. In our study, we found that the human SOX9+ REC does not express mature kidney cell markers in F-Clone culture, but had potent potential to give rise to mature tubular epithelial cells in in-vitro 3D culture system or after intra-renal transplantation. Therefore, it could have some characteristics of kidney progenitor cells. Given the large size and complexity of kidney architecture, it is very likely that multiple stem/progenitor cells co-exist in different locations of the kidney and exert distinct functions in vivo.

To investigate the human SOX9+ REC behavior in vivo, here we describe a novel, efficient approach to transplant the expanded single human SOX9+ REC into mouse kidney, to create a chimeric kidney with functional human tubules. The human SOX9+ REC derived from biopsy or urine were transplanted into the incision of kidney, which yielded large scale kidney engraftment of cells as compared to the traditional intravenous injection method which leads to limited kidney engraftment. Especially, the ability to clone nephron-regenerating SOX9+ REC from human urine, to genetically modify them, and then to successfully transplant them makes a highly useful platform for future autologous cell transplantation therapy. Of note, most of the transplanted SOX9+ REC only differentiate into proximal tubules in vivo. Human distal tubules, collecting ducts and podocytes are missing in the chimeric kidney. It seemed that the murine kidney microenvironment is biased for proximal tubule cell lineage and future work should be done to modify the cells as well as the microenvironment so that to achieve whole nephron regeneration. Altogether, our research on the special population of endogenous kidney cells improved our understanding of human kidney repair mechanisms and highlights their potential application in personalized regenerative medicine.

## Methods

### Mice, monkey and human tissue

All animal studies were performed under the guidance of Tongji University Association for Laboratory Animal Care and Use. For mouse SOX9+ REC cloning, kidneys of 8-10-week-old normal or ROSA mT/mG mice bred in SPF facility were collected to obtain renal cortex and medulla samples. For monkey SOX9+ REC cloning, the kidney of a 5-year-old male *Macaca fascicularis* was collected to obtain renal cortex, medulla and papilla samples. All mouse experiments were performed on both male and female mice and represent a minimum of n=5 mice in all groups. All mice were randomly allocated to the experimental groups.

Patients with the indicated CKD were diagnosed by K/DOQI clinical practice guidelines and all individuals had thorough medical examinations prior to sampling. For human kidney tissue sampling, percutaneous renal needle biopsies were performed to obtain patient kidney samples. Patients were in a prone position during the biopsy. Biopsies were carried out after local anesthesia with ultrasound-guided core tissue biopsy needles (18 gauge). All renal specimens were subjected to pathological diagnosis. For adult urine-derived SOX9+ REC cloning, urine samples were collected from patients and healthy volunteers. For each subject, 200ml mid-stream urine was collected into 50ml conical tubes and centrifuged to harvest cells. All the human tissues were obtained following clinical SOP under the patient’s consent and approved by the Shanghai Ruijin Hospital Ethics Committee.

### Isolation, expansion and genetic labeling of SOX9+ REC from kidney tissue and urine

For SOX9+ REC isolation from kidney tissue, samples were washed with ice-cold buffer (F12 medium containing 5% FBS and 1% Pen/Strep) and minced by sterile scalpel into 0.2-0.5 mm^3^ sizes to a viscous and homogeneous appearance. The minced tissue was then digested with dissociation buffer in room temperature overnight with gentle agitation. Digested cell suspensions were washed 5 times with wash buffer and passed through 70-μm Nylon mesh (Falcon, USA) to remove aggregates. Cell pellets were collected by centrifuge of 200×g and then seeded onto a feeder layer of lethally irradiated 3T3-J2 cells in modified SCM-6F8 medium^3^.

For SOX9+ REC isolation from human urine, the urine samples were centrifuged at 390×g for 10 minutes, the supernatant was carefully removed, and all the cell pellets from one subjects were resuspended in ice-cold urine wash buffer (F12 medium containing 5% FBS, 1% Pen/Strep, 20μg/mL Gentamicin and 2.5μg/mL Amphotericin B) and pooled into one 50ml conical tube, followed by extensive washing with urine wash buffer for 5 times. Then each cell pellet was seeded onto the feeder layer with modified SCM-6F8 medium in one well of a 12-well plate. Most urine-derived cells did not attach to the feeder layer and were removed by the first medium change on day 3.

Single cell colonies were selected for pedigree cloning using cloning cylinders and high vacuum grease. Movies of cell growth were captured using a JuLI^TM^ Stage Real-Time Cell History Recorder (NanoEnteck, Korea). For better visualization of colony growth, tdTomato+ SOX9+ REC derived from mT/mG mice were used.

For GFP labeling of cultured SOX9+ RECs, 293T cells were transfected with pLenti-CMV-eGFP plasmid together with a lentviral packaging mix (Life Technologies, USA) to produce GFP lentivirus. Lentivirus supernatant was collected, filtered and stored at -80°C until use. Medium containing lentivirus was added to the cell culture together with 10 μg/mL polybrene and incubated for 24 hours, followed by FACS sorting of single GFP+ SOX9+ REC to 96-well plates.

### Histology and immunofluorescence

For cell immunofluorescence staining, cells were fixed by 4% paraformaldehyde, and then incubated with 0.25% Triton X-100 for 8 minutes to permeabilize. For tissue histology and immunofluorescence staining, tissue samples were fixed in 3.7% formaldehyde overnight at 4°C. For cryosection, the fixed tissue was infiltrated with 30% sucrose before embedding, then embedded into the Tissue-Tek O.C.T compound (Sakura, Japan), 5–8 μm sections were prepared using a cryotome (Leica microsystem, Germany). For paraffin section, the fixed tissue was dehydrated by ethanol gradient and processed in an automatic tissue processor, and then embedded into the paraffin blocks. All the samples were sectioned at 5–10 μm thickness using a microtome (Leica microsystem, Germany). Haematoxylin and eosin (H&E) staining was performed following standard protocol. Immunofluorescence staining was conducted to reveal the protein expression in a standard protocol using the antibodies on paraffin sections, cryosections or glass smears.

Antibodies used for immunofluorescence include: SOX9 (1:200, CY5400, Abways), Krt7 (1:500, sc-53263, Santa Cruz), PAX2 (1:50, AF3364, R&D), PAX2 (1:100, ab79389, Abcam), Ki67 (1:100, 550609, BD bioscience), ATP1A1 (1:200, C464.6, Santa Cruz), SLC22A6 (1:200, ab183086, Abcam), Umod (1:200, ab207170, Abcam), CDH1 (1:500, 3195S, Cell Signaling), SLC12A1 (1:200,18970-1-AP, Proteintech), PAX8 (1:200, ab189249, Abcam), PODXL (1:200, ab150358, Abcam), SYNPO (1:200, 21064-1-AP, Proteintech), KIM-1 (1:50, AF1817, R&D), GFP (1:200, ab290, Abcam), GFP (1:500, ab6673, Abcam), tdTomato (1:200, TA180009, OriGene), HuNu (1:200, ab191181, Abcam), and human specific Lamin A+C (1:200, ab108595, Abcam). Alexa Fluor-conjugated Donkey 488/594 (1:200, Life Technologies, USA) were used as secondary antibodies. In control experiments, the primary antibodies were replaced by IgG.

### Mouse kidney injury models

To establish a UPN mouse model, 8-10-week-old C57/B6 mice were anesthetized by intraperitoneal injection of 4% chloral hydrate (0.5g/kg body weight). The left lateral peritoneum was cut to expose the left kidney and the renal artery was clamped with haemostatic forceps. About one-third of the left kidney was cut off from the upper pole of the kidney using a surgical blade. After cleaning off the blood, the incision was sealed with FuAiLe Medical glue (FAL) and the haemostatic forceps on the renal artery was removed. The muscle layer was closed with sutures (Ethicon, Germany), followed by closing of the skin.

For the UUO injury model, the abdomen was opened with a midline incision and the left kidney and upper ureter is exposed. Mice were subjected to surgical cautery of the left ureter 15 mm below the pelvis.

For wedge resection surgery model, the left lateral peritoneum was cut open to expose the left kidney and the kidney artery was clamped with small haemostatic forceps. Then a straight incision was performed on the back of the kidney using a surgical blade to make the resection. After cleaning off the blood, the kidney was sealed with FuAiLe Medical glue (FAL). Then the kidney was located back into the abdominal cavity before abdomen closing.

### Cell karyotyping

To arrest human SOX9+ REC in mitosis metaphase, cells at 75% confluence were treated with 1 μg/mL colchicines for 7 h and digested into single cells by 0.25% trypsin. Then the cells were incubated with 0.4% KCl at 37°C for 40 min and fixed by 10 mL fixation solution including methanol and glacial acetic acid (3:1) at room temperature for 30 min. The prepared cell suspension was dropped and spread on slides. Samples on slides were treated with 0.0005% trypsin for 5 min and stained with 15% Giemsa (Sigma-Aldrich, USA). Banding patterns on chromosome spreads were checked for more than 15 mitotic phases.

### Detection and calculation of somatic Copy Number Variation (CNV)

We performed analysis of the CNV from whole genome sequencing (WGS) data. Blood samples were collected from urine sample donors with signed consents and compensation. WGS was performed on genomic DNA isolated from peripheral blood mononuclear cells and the corresponding human SOX9+ RECs. All of the libraries were constructed with PCR amplification. We sequenced all sets of libraries on BGISEQ-500 at BGI TechSolutions Co. (BGI-Tech). We excluded all of the low quality raw reads, aligned the remaining high quality ones with human reference (hg19) genome. Taking as input aligned reads, Control-FREEC constructs copy number and B-allele frequency profiles. The profiles are then normalized, segmented and analyzed in order to assign genotype status (copy number and allelic content) to each genomic region. With peripheral blood mononuclear cell samples from the same individuals as control, Control-FREEC discerns somatic variants from germline ones.

### 3 dimensional culture of adult kidney and kidney organoid

*Adhered 3D culture on Matrigel.* 3D culture of cells were performed on Matrigel Matrix (Corning, USA) as previously described^5^. Cell clumps were then cultured in differentiation medium for 10 days until harvest. After lumen formation was observed for the spheres indicating proper differentiation, the cells were harvested for bulk RNA-Seq analysis.

*Non-adhered 3D culture in U-bottom plates.* Non-adhered SOX9+ REC organoid differentiation was carried out as follows. SOX9+ REC cells were harvested and seeded at 50,000 cells/well in 96-well U-bottom low cell-binding plates (S-bio, Japan) for organoid formation. The organoids were then cultured in differentiation medium for 14 days until harvest. The harvested samples were subjected to immunofluorescence, TEM, and single cell RNA-sequencing. For immunofluorescence, 5 μm cryosections were prepared and stained with tubule and glomerulus markers followed by counterstaining with DAPI.

### Transmission electron microscopy

Organoid samples were fixed in 2% glutaraldehyde (Sigma, USA) in PBS at 4°C overnight, followed by fixation with 1% osmium tetroxide in PBS at room temperature for 1.5 h.

Samples were subsequently dehydrated in a graded series of acetone, infiltrated with Epon resin (Ted Pella) in a 3:7 solution of Epon and acetone for 3 h and then a 7:3 solution of Epon and acetone overnight. Samples were then placed in 100% fresh Epon for several hours and embedded in Epon at 60 °C for 48 h. Thin sections were prepared by using the EM TXP mechanical milling system (Leica) and the EM UC6 ultramicrotome (Leica), collected on formvar-coated grids, stained with 3% uranyl acetate for 20 min and 1% lead citrate for 10 min, and examined in a JEM-1230 transmission electron microscope (Jeol) at 80 kV. Images showing glomerular structures were recorded using the Gatan798 imaging system.

### Intra-renal transplantation of SOX9+ RECs

For intra-renal transplantation, 8-10 week-old NOD/SCID mice (The Jackson Laboratory, USA) were anaesthetized, and wedge resection were performed. After cleaning off the blood and sealing cut with FuAiLe Medical glue (FAL), GFP labeled SOX9+ RECs were injected into the kidney. For each mouse, 3×10^6^ cells were used. Mice were sacrificed 1∼2 weeks after the transplantation and kidney and blood samples were harvested for further analysis.

### Dextran tracing

10,000 MW fixable Alexa Fluor-594 Dextran (Life Technologies, USA) was used to visualize vasculature 14 days after SOX9+ REC intra-renal transplantation. 200 μL 5 mg/ml Dextran was injected to mice via the tail vein. 4 hours after injection, the chimeric mice were sacrificed. The kidney samples were collected and fixed with 3.7% formaldehyde (Sigma, USA) at room temperature for 30 minutes. Samples were cryosectioned at 5 μm and observed under a fluorescence microscope (Olympus IX73, Olympus, Japan) or a confocal microscope (A1R, Nikon, Japan).

### Blood urea nitrogen (BUN) measurement

Normal mice and injured mice transplanted with saline or human RECs were sacrificed 14 days after transplantation and samples for BUN assay were collected. The measurements were conducted by using a Urea Nitrogen Colorimetric Detection Kit (Thermo Fisher Scientific, USA) following the manufacturer instruction. The amount of BUN was expressed as mg/dl.

### Isolation and culture of human-lung derived SOX9+ basal cells

The bronchoscopic procedure for lung sampling was performed by board certified respiratory physicians using a flexible fiber-optic bronchoscope (Olympus, Japan). Before the bronchoscopy, oropharyngeal and laryngeal anesthesia was obtained by administration of 2 mL of nebulized 4% lidocaine, followed by 1 mL of 2% topical lidocaine sprayed into the patient’s oral and nasal cavities. After the bronchoscope was advanced through the vocal cords, 2 mL of 2% lidocaine solution was instilled into the trachea and both main bronchi through the working channel of the bronchoscope. Then a disposable 2-mm brush was advanced through the working channel of the bronchoscope and used to collect airway epithelial cells by gently gliding the brush back and forth 2 times in 4-6 order bronchi in the right or left lobe. To isolate the human DASCs, the brush with samples were cut with scissors into 1 cm pieces. After removing sputum, the brush pieces were directly digested with dissociation buffer. Specimens were incubated at 37°C for an hour with gentle rocking. Dissociated cells were passed through 70-μm Nylon mesh and then washed twice with cold F12 medium. Cell were plated onto feeder cells in DASC culture medium for lung including DMEM/F12, 10% FBS (Hyclone, Australia), Pen/Strep, amphotericin and growth factor cocktail as previously described^5^. Under 7.5% CO2 culture condition, the DASC colonies emerged 3–5 days after plating, and were digested by 0.25% trypsin-EDTA (Gibco, USA) for 3–5 min for passaging. Typically, DASCs are passaged every 5 to 7 days and split at 1:7 ratio.

### Microarray analysis, bulk RNA-sequencing analysis and bioinformatics

Mouse tissues, SOX9+ REC and organoids were characterized by transcriptome microarray (Affymetrix, USA) to study their gene expression profiles. Duplicate experiments for microarray were taken from 2 biological samples. Briefly, total RNA was checked for a RIN number to inspect RNA integrity by an Agilent Bioanalyzer 2100 (Agilent technologies, Santa Clara, CA, US). Double-stranded cDNA from samples were synthesized, labeled and hybridized to Mouse Genome 430 2.0 array (Affymetrix, Santa Clara, CA, USA) and the chips were scanned by GeneChip® Scanner 3000 (Cat#00-00212, Affymetrix, Santa Clara, CA, US) and Command Console Software 4.0 (Affymetrix, Santa Clara, CA, US) with default settings. Raw data were normalized by MAS 5.0 algorithm, Affy packages in R.

Human kidney tissue, SOX9+ RECs and 3D cultured spheres were analyzed by bulk RNA-sequencing. Duplicate experiments were taken from 2 biological samples. RNA was extracted using RNeasy® Mini Kit(REF# 74104) and a total amount of 3 μg RNA per sample was used as input material for the RNA sample preparations. Sequencing libraries were generated using NEBNext® UltraTM RNA Library Prep Kit for Illumina® (NEB, USA) following manufacturer’s recommendations and index codes were added to attribute sequences to each sample. Raw data (raw reads) of FASTQ format were firstly processed through in-house PERL scripts. In this step, clean data (clean reads) were obtained by removing reads containing adapter, reads containing ploy-N and low quality reads from raw data. All the downstream analyses were based on the clean data with high quality. FASTQ files were aligned against the human reference (hg19/hGRC38) genome. Index of the reference genome was built using STAR (v2.5.1b) and paired-end clean reads were aligned to the reference genome. HTSeq v0.6.0 was used to count the reads numbers mapped to each gene, and then FPKM of each gene was calculated based on the length of the gene and read counts mapped to this gene. Differential expression analysis of two conditions/groups (two biological replicates per condition) was performed using the DESeq2 R package (1.10.1). The resulting P-values were adjusted using the Benjamini and Hochberg’s approach for controlling the false discovery rate. Genes with an adjusted P-value <0.05 found by DESeq2 were assigned as differentially expressed. Corrected P-value of 0.05 and an absolute fold change of 2 were set as the threshold for significant differential expression. Heatmap and volcano plot were generated using R scripts. PPI analysis of differentially expressed genes was visualized with Cytoscape (v3.2.0) and based on the STRING database. The program “prcomp”, a built-in program for principal component analysis (PCA) in R packages, was employed with default parameters. We evaluated the variance percentage of each principal component, and found the top 3 components accounted for 83.9% of the total variance, where PC1 accounted for 49.5%, PC2 23.2% and PC3 11.2%. Another program “ggplot2” (v3.1.0) in the R packages was used to plot the 2D view of PCA. Gene Ontology (GO) enrichment analysis of differentially expressed genes was implemented by the cluster Profiler R package, in which gene length bias was corrected. GO terms with corrected P value less than 0.05 were considered significantly enriched by differentially expressed genes.

### Single cell RNA sequencing analysis

Single cells were captured and barcoded in 10x Chromium Controller (10x Genomics). Subsequently, RNA from the barcoded cells were reverse-transcribed and sequencing libraries were prepared using Chromium Single Cell 3’v3 Reagent Kit (10x Genomics) according to manufacturer’s instructions. Sequencing libraries were loaded on an Illumina NovaSeq with 2×150 paired-end kits at Novogene, China. Raw sequencing reads were processed using the Cell Ranger v.3.1.0 pipeline from 10X Genomics. In brief, reads were demultiplexed, aligned to the human GRCh38 genome and UMI counts were quantified per gene per cell to generate a gene-barcode matrix. Data were aggregated and normalized to the same sequencing depth, resulting in a combined gene-barcode matrix of all samples. Seurat v.3.0 was used for quality control, dimensionality reduction and cell clustering. We removed the low-quality cells with less than 200 or more than 9000 detected genes, or if their mitochondrial gene content was > 10%. Genes were filtered out that were detected in less than 3 cells. This filtering step resulted in 20,205 genes X 5098 cells. We used the function SCTransform to normalize the data which can replace three functions NormalizeData, ScaleData, and FindVariableFeatures. Before clustering, variants arising from number of genes, number of UMIs, and percentage of mitochondrial reads were regressed out by specifying the vars.to.regress argument in Seurat function SCTransform. We performed principal component analysis (PCA) dimensionality reduction with the highly variable genes as input in Seurat function RunPCA. We then selected the top 30 significant PCs for two dimensional t-distributed stochastic neighbor embedding (tSNE), implemented by the Seurat software with the default parameters. We used FindCluster in Seurat to identify cell clusters. To identify the marker genes, differential expression analysis was performed by the function FindAllMarkers in Seurat with likelihood-ratio test. Differentially expressed genes that were expressed at least in 25% cells within the cluster and with a fold change more than 0.25 (log scale) were considered to be marker genes. Featureplots were performed by the function Featureplot in Seurat with the default parameters. Threshold of legends were adjusted with the R package ggpolt2.

### Quantitative Real Time-PCR

RNA was isolated from snap-frozen kidney tissues or SOX9+ REC stored at -80°C using an Rneasy Mini kit (QIAGEN) according to the manufacturer’s instruction. All RNA was digested with DNase I (Takara, Japan). 1 ug total RNA and PrimeScript RT Master Mix (Takara, Japan) was used for reverse transcription in a SimpliAmp Thermal Cycler (Life technologies, USA). qRT-PCR was performed in triplicate using an ABI7500 Sequence Detection System (Applied Biosystems, USA) and SYBR Premix Ex Taq II (Takara, Japan).

DNA primer pairs were designed to span exons, when possible, to ensure that the product was from mRNA. The following cycling conditions were used: 1 cycle of 50°C for 2 minutes, 1 cycle of 90°C for 5 minutes and 40 cycles of 95°C for 12 seconds and 60° for 1 minute. The specificity of the amplified product was evaluated using the melting curve analysis. Internal control glyceraldehyde 3-phosphate dehydrogenase (GAPDH) was used to normalize the result in each reaction, and relative fold change was calculated by 2^-ΔΔCt^ method. The following primer pairs were used: for human AQP1: 5’-CCCGAGTTCACACCATCAG-3’ and 5’-CTCATGTACATCATCGCCCA-3’, for human CDH1: 5’-GACCGGTGCAATCTTCAAA-3’ and 5’-TTGACGCCGAGAGCTACAC-3’, for human NPHS2: 5’-TGAGGACAAGAAGCCACTCA-3’ and 5’-ACGTGGATGAGGTCCGAG-3’, for human GAPDH: 5’-AGTATGACAACAGCCTCAAGAT-3’ and 5’-GTCCTTCCACGATACCAAA-3’.

### Cisplatin nephrotoxicity assay

8-10-week-old NOD-SCID mice maintained in SPF animal facilities were used for human DASC and human REC double transplantation. Mouse lung was preconditioned by intratracheally instilling bleomycin (Selleckchem, USA) at 3 U/kg body weight 8 days prior to human DASC transplantation. On the day of DASC transplantation, mice were anesthetized using 3.7% chloral hydrate and 1×10^6^ eGFP labeled human DASCs were suspended in 50 μL PBS and transplanted to each mouse. Intratracheal aspiration was performed by injecting the cells into the airway. On the same day mice were subjected to human REC transplantation as described above to generate dual transplanted mice. 7 days after cell transplantation, nephrotoxicity model was achieved by subcutaneous injection of the renal tubule specific toxic drug cis-diaminedichloroplatinum (cisplatin) at the dose of 10 mg/kg. 3 days after cisplatin treatment, mice lungs and kidneys were harvested for immunofluorescence staining and Annexin V apoptosis assay.

### Flow cytometric analysis of apoptosis

For Annexin V apoptosis assay, harvested lungs and kidneys were digested into single cell suspensions, washed 5 times with ice-cold PBS and then resuspended in 1× binding buffer (BD Biosciences, USA) at a concentration of 1 × 10^6^ cells/mL. 100 μL of the cell suspension, 5 μL of APC-conjugated Annexin V (BD Biosciences, USA) and 5 μL of PI (BD Biosciences, USA) were sequentially added to a FACS tube followed by gently shaking and incubation for 15 min at room temperature in the dark. Following incubation, 400 μL of 1× binding buffer was added to each tube and cells were analyzed on a BD FACS VerseTM system within 1 hour. Gate was defined to remove debris and doublet cells using FSC and SSC. Transplanted human cells were identified by detection of GFP signaling. Live and dead GFP positive cells were separated by APC-conjugated Annexin V and PI.

### Statistical analysis

Block randomization was used to randomize mice and samples into groups of similar sample size. No data was intentionally excluded. Postoperative deaths of animals were excluded from study. Statistical power analysis was used to ensure adequate sample size for detecting significant differences between samples. All experiments were assessed by at least two blinded participating investigators. Results are presented as means ± SD.

GraphPad Prism (version 7.0a) or R programming were used for data management, statistical analysis and graph generation. Matched results were assessed using Wilcoxon matched-pairs signed rank test or matched two-way ANOVA according to normality. Comparisons among multiple groups were analyzed using Tukey’s multiple comparison test. The exact number (n) of sample size for each experiment was stated in the corresponding figure legend. Differences with P ≤ 0.05 were considered statistically significant.

---

**Supplementary Figure 1.**
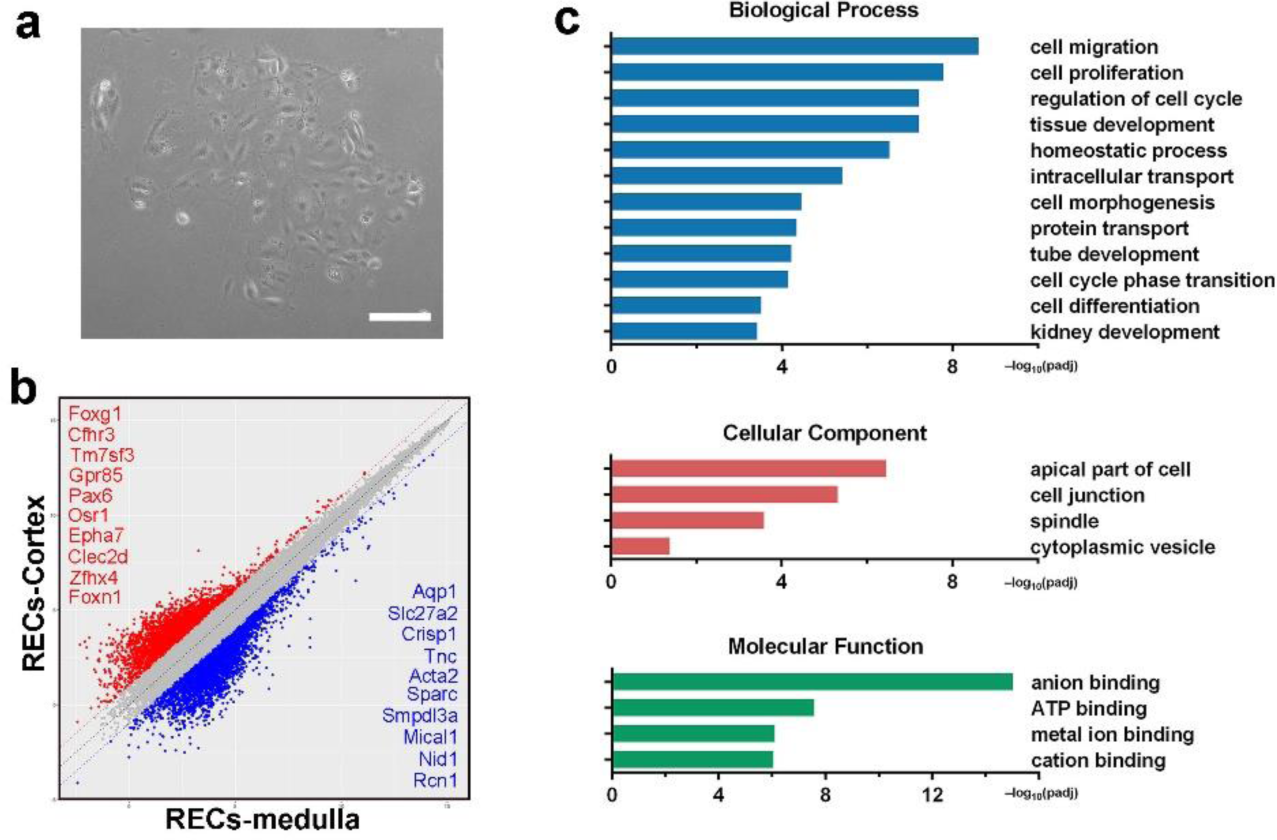
Mouse cortex and medulla-derived REC colonies. (**a**) Representative images of REC culture using conventional culturing method demonstrating slow cell growth and senescence-like cell morphology. Tissues from normal adult mouse kidneys were dissected and digested to single cell suspensions. Regular DMEM containing 10% (v/v) fetal bovine serum was used as culture medium. Scale bar, 50 μm. (**b**) Scatter plot showing the gene expression pattern mouse cortex and medullar derived Sox9+ REC with top ranked marker genes listed, respectively. (**c**) Gene ontology enrichment analysis of differentially expressed genes in medulla derived Sox9+ REC versus mouse medulla tissue.

**Supplementary Figure 2.**
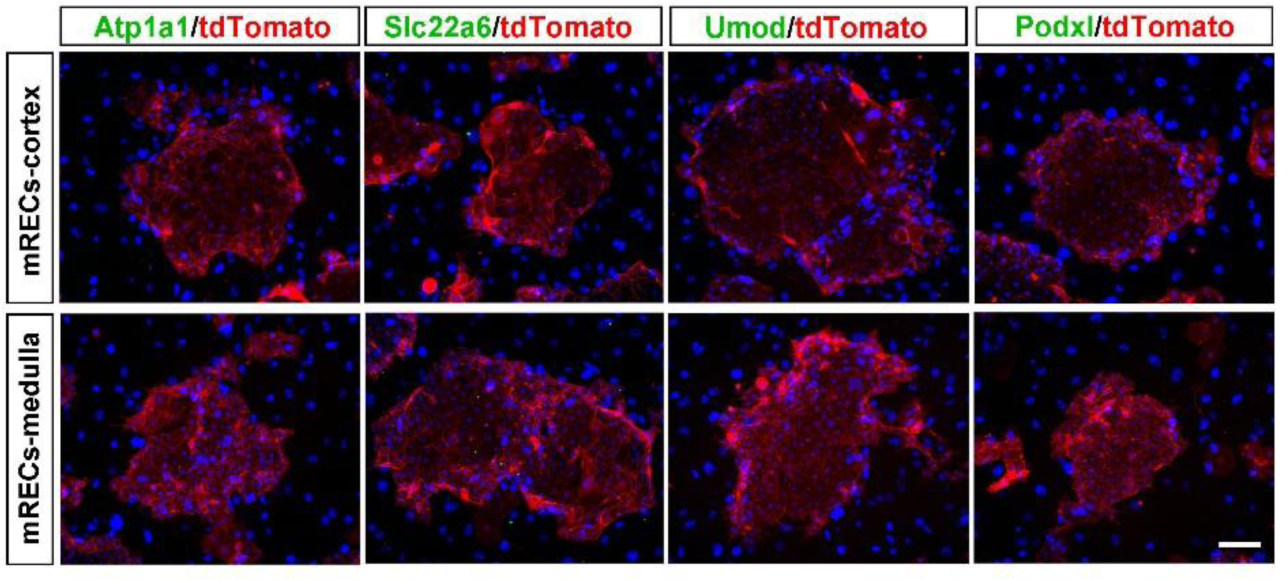
Tubular and glomerulus marker expression in mouse cortex and medulla-derived REC colonies. Immunostaining of Sox9+ REC colonies derived from the cortex or medulla with antibodies against mature tubular marker Atp1a1, Slc22a6, Umod, and mature glomerulus marker Podxl. Scale bar, 50 μm.

**Supplementary Figure 3.**
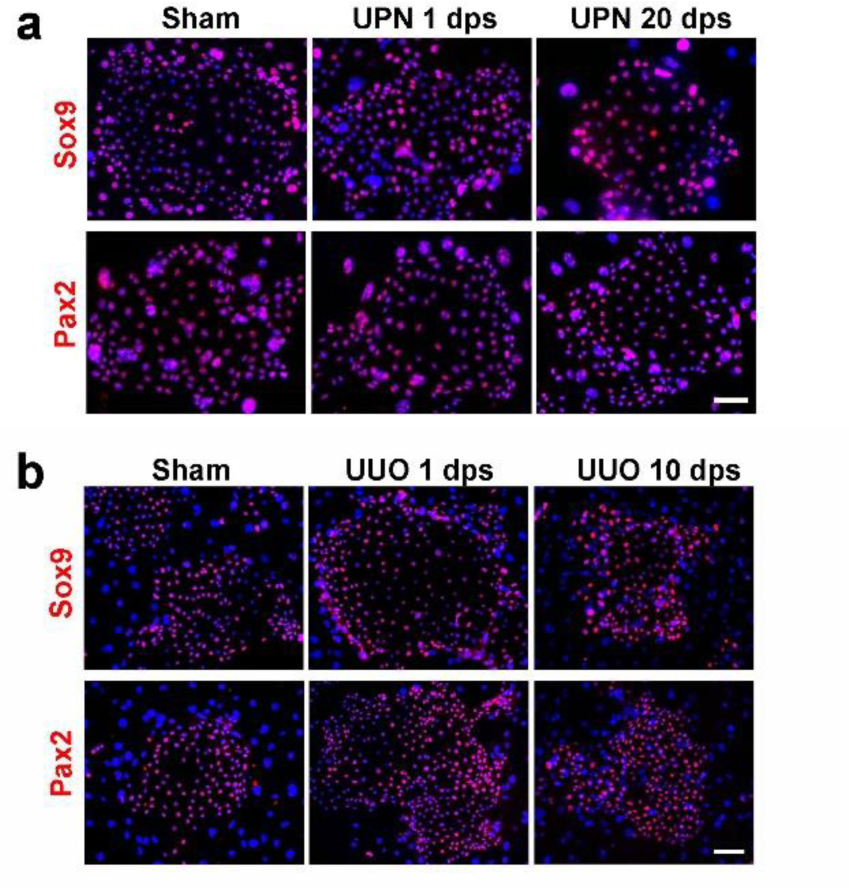
Characterization of Sox9+ REC colonies derived from UPN and UUO injured kidneys. (**a**) Sox9+ REC colonies derived from sham and UPN injured kidneys stained with Sox9 and Pax2 antibodies. Scale bar, 50 μm. (**b**) Sox9+ REC colonies derived from sham and UUO injured kidneys stained with Sox9 and Pax2 antibodies. Scale bar, 50 μm.

**Supplementary Figure 4.**
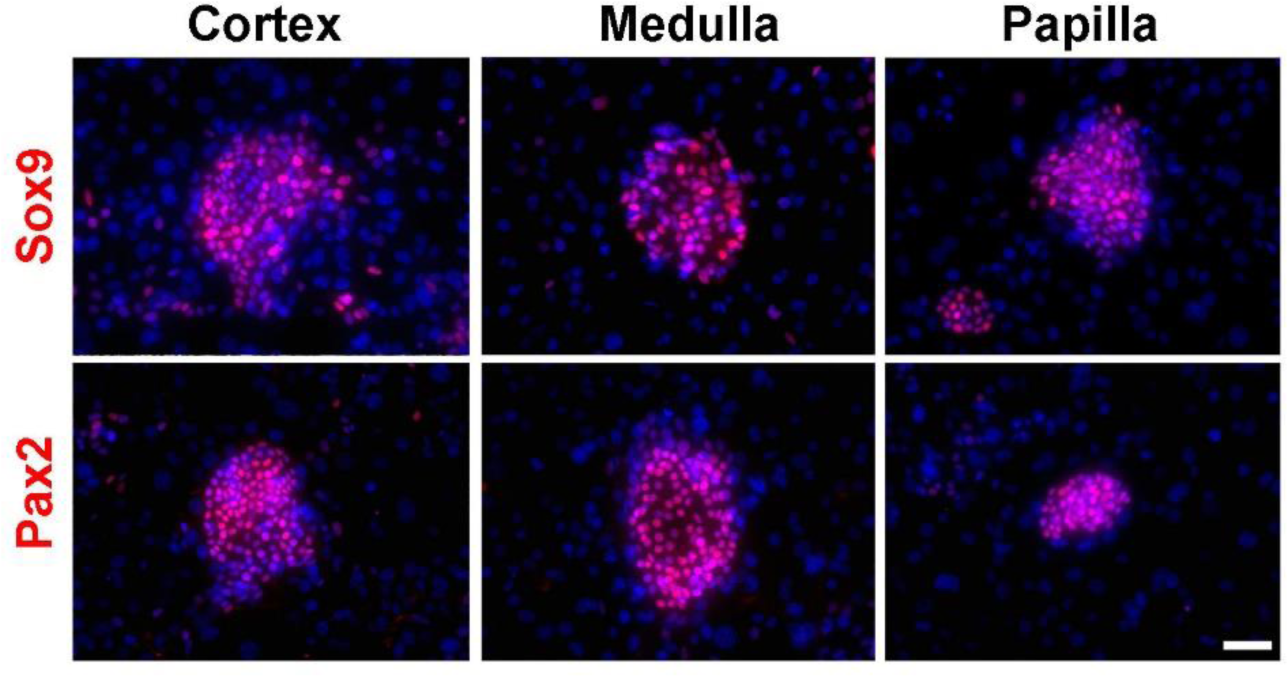
Characterization of Sox9+ REC colonies derived from monkey kidneys. Immunofluorescence staining of monkey cortex-, medulla-, and papilla-derived Sox9+ REC colonies. Scale bar, 50 μm.

**Supplementary Figure 5.**
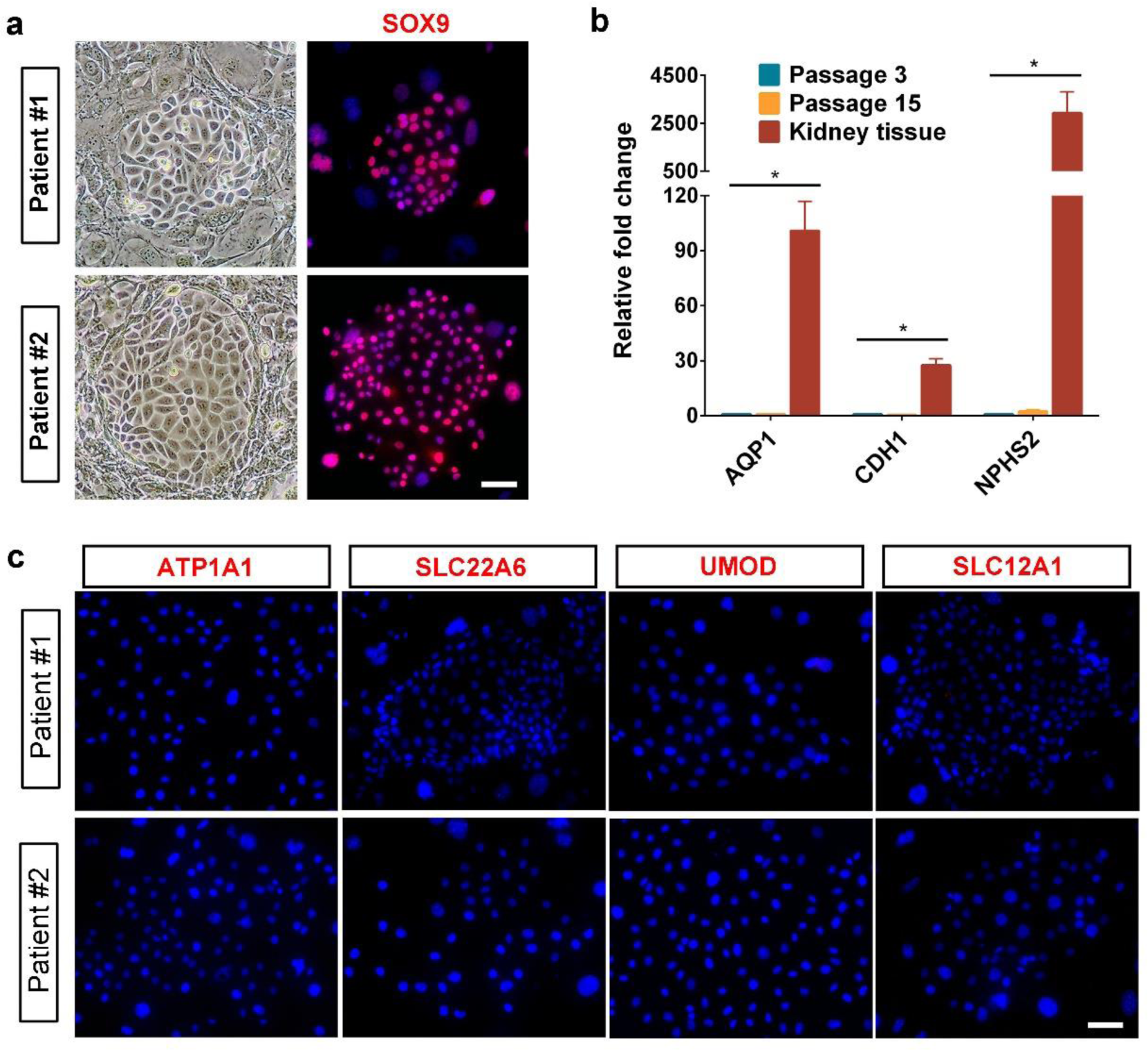
Human SOX9+ REC colonies derived from kidneys of patients with CKD. (**a**) Human SOX9+ REC colonies isolated from needle renal biopsy of two patients stained with progenitor cell marker SOX9. Representative image of n=2 independent experiments. Scale bar, 20 μm. (**b**) qRT-PCR showing relative expression of mature kidney markers in early and late passage SOX9+ REC and kidney tissue. Analysis were performed in triplicate for n=3 independent qRT-PCR experiments. Statistics are inclusive of all biological replicates. *p < 0.05. (**c**) Human REC colonies from kidneys of two patients stained with indicated mature renal cell markers showed no expression at all. Scale bar, 20 μm.

**Supplementary Figure 6.**
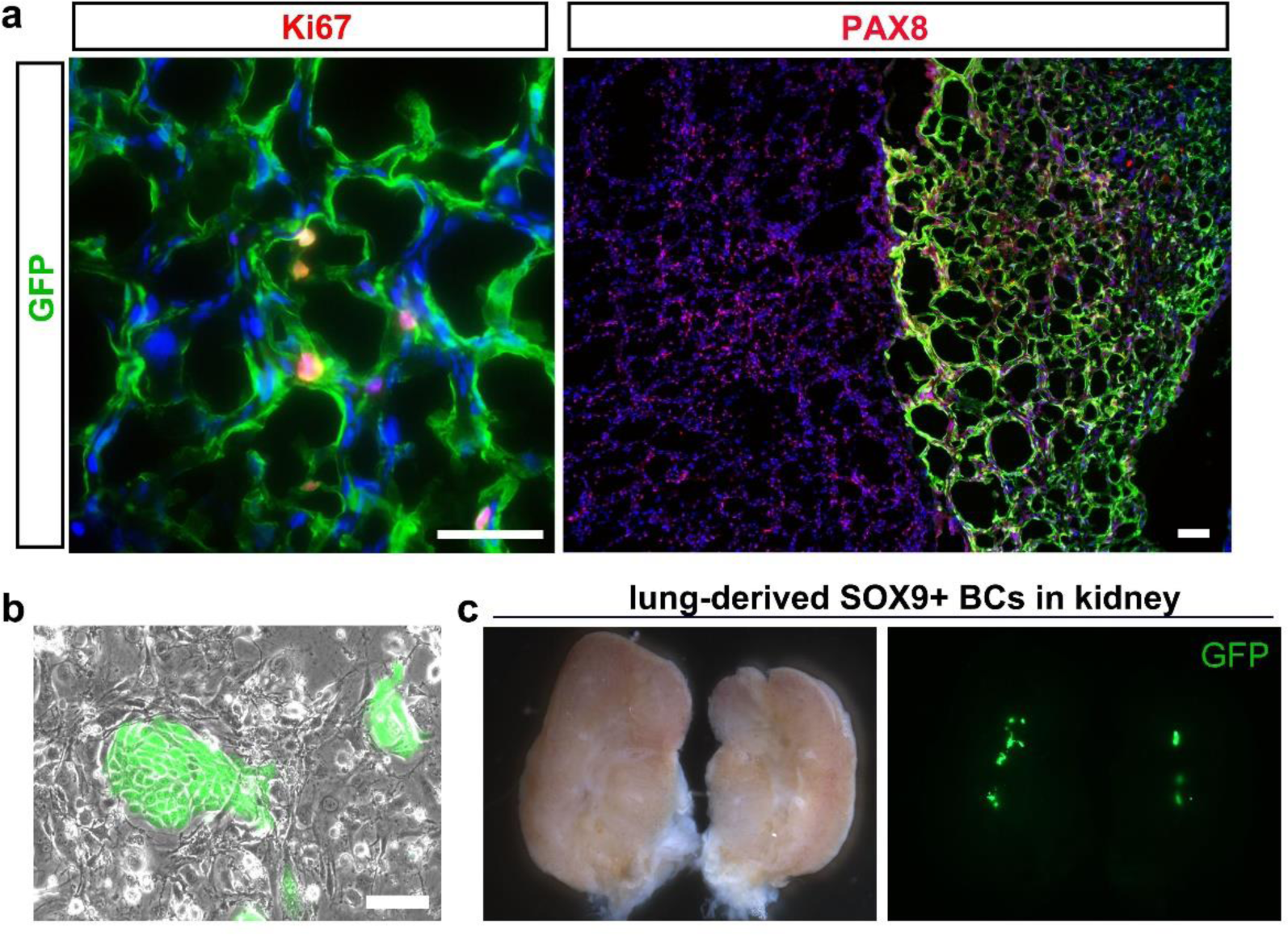
Intra-renal transplantation of organ-specific cells. (**a**) Immunostaining on sections of single cell-derived SOX9+ REC transplanted area 14 days after transplantation with PAX8 (tubular epithelium) and Ki67 (proliferative cell) antibodies. Scale bars, 50 μm. (**b**) Representative image of GFP-labeled human DASC colonies. Scale bars, 20 μm. (**c**) Bright field and direct fluorescence imaging of mouse kidney intrarenally transplanted with GFP-labeled human lung-derived SOX9+ basal cells showing minimal engraftment. BCs, basal cells. Representative of n=3 independent experiments.

**Supplementary Table 1.** Clinical data of CKD patients and healthy donors.

| Patient | Age<br>(Years) | Gender | Sample | Group | Diagnosis | Success in<br>achieving<br>SOX9+<br>REC culture |
| --- | --- | --- | --- | --- | --- | --- |
| 1# | 29 | F | Kidney biopsy | Patient | Tubulointerstitial nephritis | Yes |
| 2# | 60 | M | Kidney biopsy | Patient | ANCA-associated renal<br>vasculitis | Yes |
| 3# | 22 | F | Kidney biopsy | Patient | Membranous nephropathy | Yes |
| 4# | 38 | M | Urine | Patient | CKDV | Yes |
| 5# | 32 | M | Urine | Patient | CKDII | Yes |
| 6# | 16 | M | Urine | Patient | Hemoglobinuria | Yes |
| 7# | 41 | M | Urine | Patient | Hypertensive nephropathy | No |
| 8# | 57 | M | Urine | Patient | Diabetic nephropathy | No |
| 9# | 44 | M | Urine | Patient | CKDIII, hyperuricemia | Yes |
| 10# | 38 | M | Urine | Patient | CKDIII | Yes |
| 11# | 39 | M | Urine | Healthy | / | No |
| 12# | 24 | M | Urine | Healthy | / | Yes |
| 13# | 27 | F | Urine | Healthy | / | Yes |

